# HCV E1 influences the fitness landscape of E2 and may enhance escape from E2-specific antibodies

**DOI:** 10.1101/2023.04.03.535505

**Authors:** Hang Zhang, Rowena A. Bull, Ahmed A. Quadeer, Matthew R. McKay

## Abstract

The Hepatitis C virus (HCV) envelope glycoprotein E1 forms a noncovalent heterodimer with E2, the main target of neutralizing antibodies. How E1-E2 interactions influence viral fitness and contribute to resistance to E2-specific antibodies remains largely unknown. We investigate this problem using a combination of fitness landscape and evolutionary modelling. Our analysis indicates that E1 and E2 proteins collectively mediate viral fitness, and suggests that fitness-compensating E1 mutations may accelerate escape from E2-targeting antibodies. Our analysis also identifies a set of E2-specific human monoclonal antibodies that are predicted to be especially resilient to escape via genetic variation in both E1 and E2, providing directions for robust HCV vaccine development.

## I. Introduction

Hepatitis C virus (HCV), a single-stranded RNA virus, is the major cause of liver-associated disease and liver cancer. Currently, an estimated 58 million people are chronically infected with HCV [1]. Although direct-acting antivirals (DAAs) have been developed and offer promising treatments for chronic HCV infections, their high cost and low rates of HCV diagnosis limit their accessibility to a subset of infected individuals only [1], [2]. Additionally, the efficacy of DAAs is limited by their inability to prevent reinfection and the emergence of drug-resistant viral strains [3], [4]. Therefore, developing an effective vaccine is crucial for eradication of HCV.

HCV encodes a single polyprotein, which is further cleaved by cellular and viral proteases into three structural proteins (core, E1, and E2) and seven non-structural proteins (NS1, NS2, NS3, NS4A, NS4B, NS5A, and NS5B). The envelope protein E2 is vital for viral entry into liver cells (hepatocytes), and it is the major target of neutralizing antibodies elicited against HCV [5]. Studies have shown that E2 alone can induce a potent humoral immune response and serve as a promising vaccine candidate [6]–[8]. However, the other envelope protein E1, which forms noncovalent heterodimers with E2 [5], has a function that has been shown to be inter-dependent with E2 [9]–[14]. For instance, E1 helps E2 maintain its functional conformation and regulates E2’s interaction with HCV receptors CD81 and SR-B1 [10]–[13], and both E1 and E2 are needed for interaction with CLDN1 [13], a key factor in HCV entry. E1 has also been shown to modulate the folding of E2 [15], [16]. While preliminary experiments suggest that specific mutations in E1 and E2 may jointly modulate viral infectivity [13], a comprehensive analysis of the role of E1E2 inter-protein interactions in mediating viral fitness is still lacking. Moreover, fitness of HCV is closely related to its ability to escape from antibody responses [17], [18]. Therefore, investigating the effect of E1 on escape from E2-specific neutralizing antibodies is of particular interest.

To study the role of E1E2 inter-protein interactions in mediating viral fitness and antibody evasion, we develop a computational fitness landscape model, the joint model (JM), that considers interactions between E1 and E2 proteins. Comparing JM with a model that considers E1 and E2 proteins independently, named the independent model (IM), we find that JM captures more correlated structure in the E1E2 protein sequence data, providing a statistical quantification of the mutational interactions between E1 and E2. Comparing with the available in-vitro infectivity data, JM is found to be a better representative of the intrinsic fitness landscape of E1E2 compared to IM. These results suggest that E1 and E2 proteins collectively mediate infectivity of the virus. Based on the JM, we find that the strong E1E2 inter-protein interactions might be compensatory instead of antagonistic.

To investigate the role of E1 on E2 in antibody evasion, we incorporate the JM into an in-host evolutionary model and assess antibody escape dynamics. We predict that the residues with known escape mutations would be easier to escape according to the JM compared with an E2-only model [19]. This points to the potential role of E1 in facilitating escape from E2-targeting antibodies. We use the evolutionary model to study the efficacy of human monoclonal antibodies (HmAbs) known for HCV, and compare the results with those obtained from the E2-only model. We find, again, that E1 may facilitate the virus to escape specific HmAbs targeting E2. Our analysis also reveals potentially escape-resistant HmAbs against the E1E2 complex, offering directions for the development of an effective vaccine against HCV.

## II. Results

### A. Inference and statistical validation of the joint model for the E1E2 protein

We developed a computational model, termed JM, for the entire E1E2 protein using the sequence data available for subtype 1a. This model uses a maximum entropy approach to estimate the probability of observing a virus with a specific E1E2 protein sequence. In this model, the probability of any sequence **x** = [*x*_1_*, x*_2_*,..., x_N_*] is given by

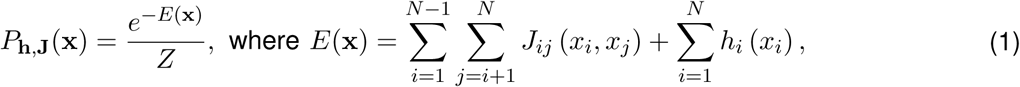

where *N* is the length of the sequence, and *Z* = Σ_x_ *e*^−*E*_h,J_(x)^ is a normalization factor which ensures the probabilities sum to one. The fields **h** and couplings **J** parameters represent the effect of mutations at a single residue and interactions between mutations at two different residues, respectively. *E*(**x**) denotes the energy of sequence **x**, which is inversely related to its prevalence. Inference of a maximum entropy model involves choosing the fields and couplings such that the model can reproduce the single and double mutant probabilities observed in the E1E2 sequence data.

We inferred the E1E2 maximum entropy model using the GUI-based software implementation of MPF-BML [20] (see Methods for details), an efficient inference framework introduced in [21]. The single and double mutant probabilities obtained from the JM matched well with the E1E2 sequence data (Fig. 1a, b). Although not explicitly included in model inference, additional statistics including the connected correlations and the distribution of the number of mutations computed from the model also agreed well with those obtained from the E1E2 sequence data (Fig. 1c, d), demonstrating the predictive power of the inferred model. Overall, these results indicate that the inferred E1E2 JM captures well the statistics of the data.

**Fig. 1:**
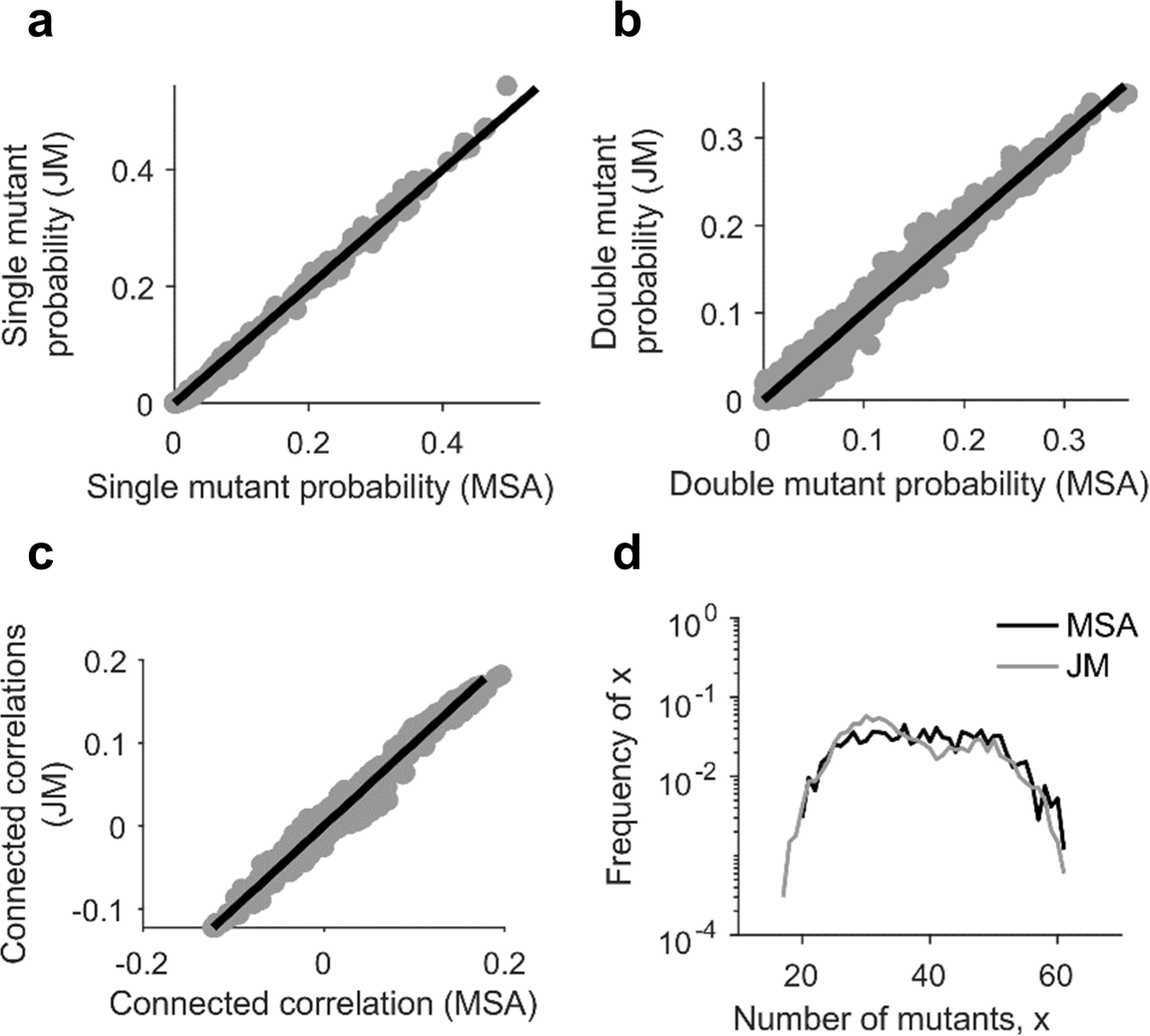
Statistical validation of the inferred E1E2 JM. Comparison of the (**a**) single mutant probabilities, (**b**) double mutant probabilities, (**c**) connected correlations, and (**d**) distribution of the number of mutants per sequence obtained from the MSA and those predicted by the inferred JM. Samples were generated from the inferred model using the MCMC method [25].

### B. E1E2 inter-protein interactions are important in mediating viral fitness

While some studies have considered E2 alone (i.e., independent of E1) [6]–[8], multiple studies have reported that these two proteins are functionally inter-dependent [10]–[13]. This suggests that interactions between E1 and E2 may be critical. Previously, we had investigated E2 alone wherein we had inferred a fitness landscape model for E2 and used it to explore HCV escape dynamics from neutralizing antibodies [19], [22]. Here, to investigate the importance of E1E2 inter-protein interactions on virus fitness and immune escape, we compared the inferred JM with a model that considers E1 and E2 proteins to be independent (see Methods for details). We refer to it as the independent model (IM). In this model, the energy of an E1E2 protein sequence x = [x_E1_, x_E2_] is given by the sum of the energies of its E1 and E2 parts, x_E1_ and x_E2_, respectively:

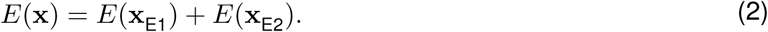

Here, *E*(x_E1_) and *E*(x_E2_) are computed separately using inferred E1-only and E2-only maximum entropy models, respectively (see Methods for details). Both the E1-only and E2-only models capture well the statistics of the respective sequence data (Supplementary Fig. S1).

Equipped with the JM and IM models, we first investigated whether E1 and E2 proteins can be considered statistically independent. This can be quantified by comparing the fraction of the correlated structure (FCS) of the E1E2 protein complex captured by the two models [23]. FCS captured by a model can be estimated using the entropy of the synthetic sequence data generated by the inferred model and comparing it with the entropy of an independent model and the estimated true entropy of the data (see Methods for details). If FCSs captured by both the JM and IM models are similar, it will be suggestive of E1 and E2 to be independent. Based on our analysis, the average FCS of the E1E2 protein complex captured by the JM (63%) was 22% more than that captured by the IM (41%) (*p* = 9.1 × 10^−5^; Fig. 2), suggesting that E1 and E2 proteins are not statistically independent. Thus, there seem to be significant inter-protein correlations that are not captured by the IM.

**Fig. 2:**
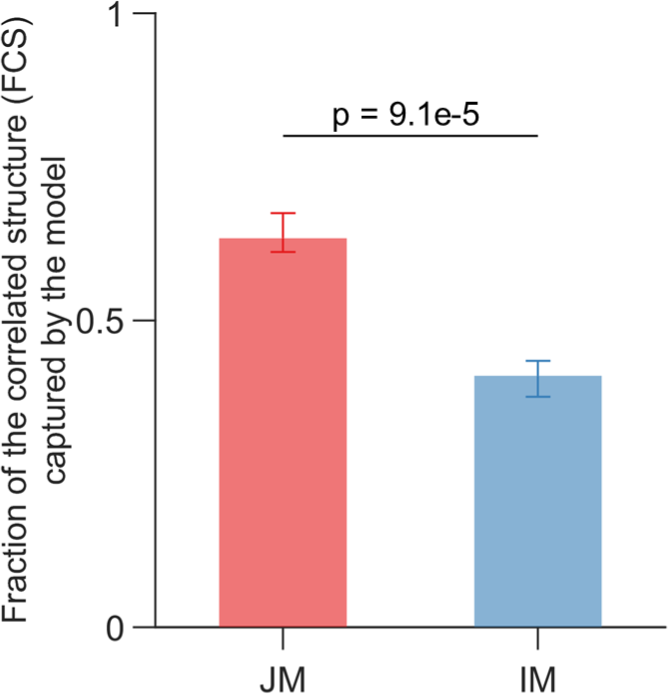
Comparison of the fraction of the correlated structure in E1E2 protein captured by the joint model (JM) and the independent model (IM) Fraction of the correlated structure (FCS) captured by a model is quantified by *I*_model_*/I*. Here, *I* = *S*_ind_ *S*_true_ is the multi-information which measures the overall strength of correlations in the system, where *S*_ind_ denotes the entropy of a site-independent model of E1E2 protein and *S*_true_ is the true entropy of the E1E2 complex estimated using the approach in [81] (see Methods for details). Similar to *I*, *I*_model_ = *S*_ind_ *S*_model_ measures the strength of correlations captured by the JM or IM model, where *S*_model_ is the entropy predicted by the JM or IM model based on the data generated using the MCMC method (see Methods for details) [23]. Entropies for the JM and IM were calculated over 10 instances of MCMC runs, and the *p*-value was calculated using the one-sided Mann-Whitney test.

We next investigated if the additional correlations captured by the JM, compared to IM, make it a better representative of the intrinsic E1E2 fitness landscape. Maximum entropy models have been shown previously to be good representatives of the underlying fitness landscapes for multiple individual viral proteins of HCV (polymerase [24] and E2 [19], [22]) and HIV [21], [25]–[28]. To test this for JM and IM, we compared the predictions of both models using the in-vitro infectivity measurements available for E1E2. We compiled a total of 156 in-vitro infectivity measurements for E1E2 from 16 studies [13], [29]–[43]. We found that the JM provided a stronger negative Spearman correlation (*r* = −0.70; see Methods for details) between the predicted sequence energies (inversely related to prevalence) and experimental fitness values (Fig. 3) than the IM (*r* = −0.54; Fig. 3, inset). This result suggests that JM is a better representative of the E1E2 fitness landscape. It also indicates the potential importance of E1E2 inter-protein interactions in mediating viral fitness.

**Fig. 3:**
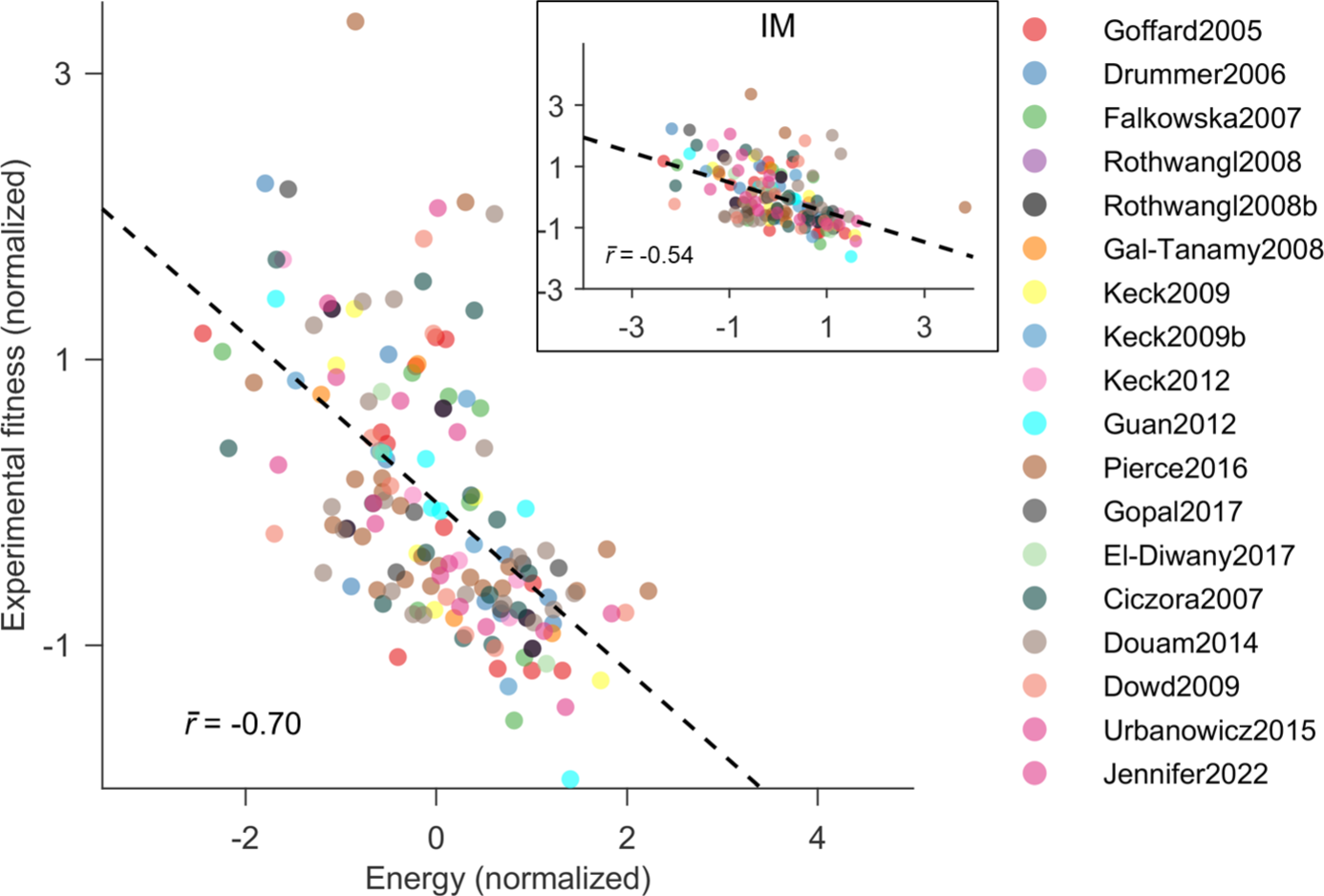
Comparison of the E1E2 fitness prediction by JM and IM. Normalized energies computed from the inferred JM correlate strongly with the experimental fitness measurements. Conversely, the inferred IM provided a much lower correlation (inset). Legend shows the references from which fitness/infectivity measurements were compiled [13], [29]–[43].

### C. Majority of strong E1E2 inter-protein interactions are compensatory

The couplings of the inferred maximum entropy model (*J_ij_* in Eq. 1) are informative of the type of interactions between residues [44], [45]. When the value of *J_ij_* is large and positive, it signifies a strong antagonistic interaction or negative epistasis between residues *i* and *j*. This results in a decrease in the fitness of double mutants and makes it harder for new mutations to occur [46]. On the other hand, when the value of *J_ij_* is large and negative in Eq. 1, it indicates a strong compensatory interaction or positive epistasis between residues *i* and *j*. This signifies improved replication of double mutants, allowing the virus to acquire diverse mutations.

Analyzing the top 300 pairs of inter-protein couplings (listed in Supplementary Data 1), i.e., with large absolute values of *J_ij_*, we found that the majority (70%) were negative (Fig. 4). This suggests that the top inter-protein couplings are largely compensatory, and that simultaneous mutations in the two proteins may assist in maintaining a viable virus. This result was robust to the number of top inter-protein couplings considered (Supplementary Fig. S2). A recent study reported E1E2 as a highly fragile complex, with 92% of alanine mutations introduced independently at each residue severely impacting virus fitness [43]. The strong compensatory interactions identified in our analysis indicate a potential mechanism by which E1 and E2, the most variable HCV proteins, may make multiple mutations while maintaining viral fitness.

**Fig. 4:**
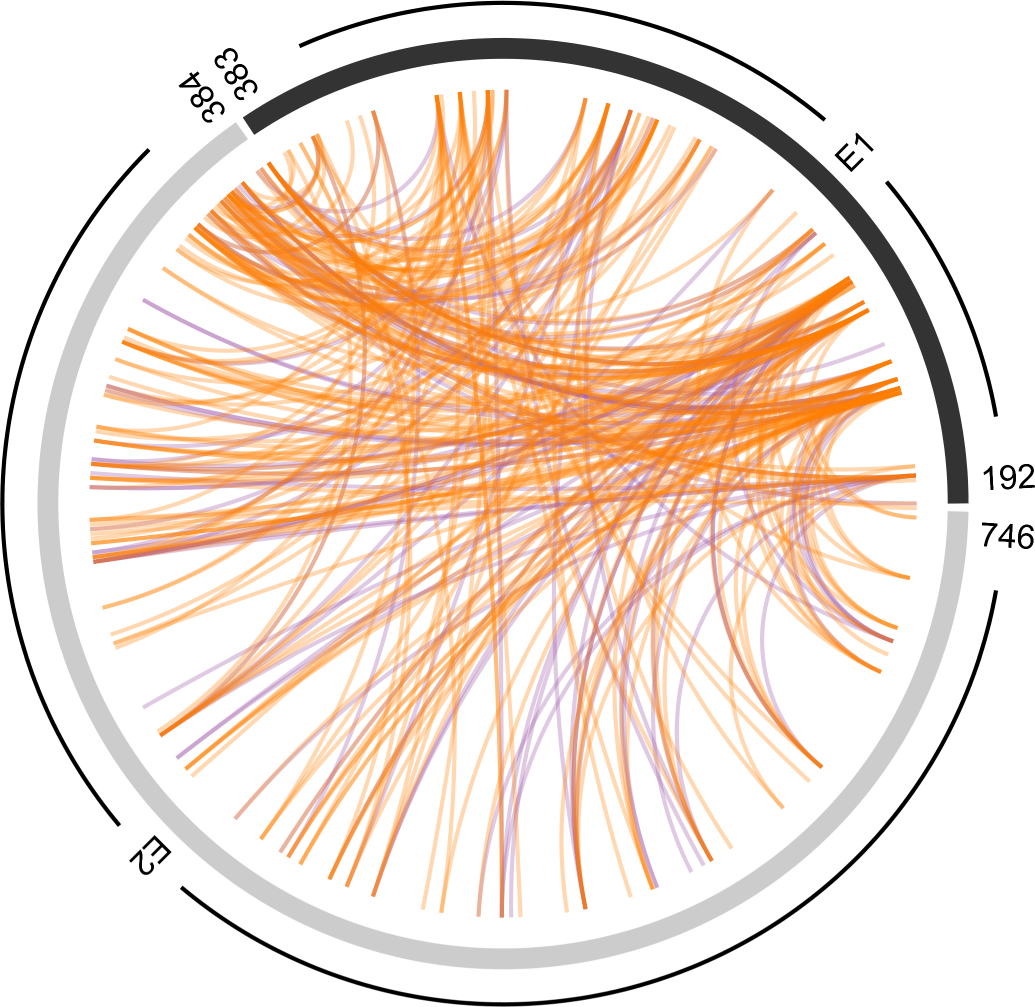
Strong E1E2 inter-protein interactions are largely compensatory. Each pair of mutations between E1 and E2 proteins was ranked by the absolute values of *J_ij_* from Eq. 1 and top 300 pairs are plotted here. Compensatory interactions (negative values of *J_ij_*) are colored in orange, while antagonistic interactions (positive values of *J_ij_*) are colored in purple. The outer segments of the circle represent E1 (shown in black, encompassing residues 192-383) and E2 (shown in gray, encompassing residues 384-746) proteins.

We further quantified whether the strongly-coupled residues (those associated with top 300 pairs of inter-protein couplings) were enriched in any known functional region of E1 and E2 proteins (see Methods for details). We observed that the strongly-coupled residues were statistically significantly enriched in hypervariable region 1 (HVR1) and hypervariable region 2 (HVR2) of E2 (Supplementary Table S1), suggesting that these regions may be involved in interactions with E1. This is also consistent with the literature that has shown that HVR2 is essential for the formation of the E1E2 heterodimer [47], and epistatic interactions exist between E1 and HVR1 of E2 [48]. As their names suggest, these two regions are highly variable, and are known to modulate viral escape from neutralizing antibodies [49]. Hence, the potential compensatory interactions between E1 and these two E2 regions may contribute to viral immune evasion.

### D. Evolutionary simulations suggest the E1 protein contributes to escape from E2-specific antibody responses

To gain a deeper understanding of the impact of E1 on viral escape dynamics from E2-specific antibody responses, we quantified and compared the average time it takes for E2 residues to escape with and without the influence of E1. To achieve this, we utilized an in-host evolutionary model that takes into account the stochastic dynamics of viral evolution within the host including virus-host interactions, virus-virus competition, and escape pathways that the virus may employ to evade immune pressure. Similar models have been used previously for simulating in-host viral evolution for HIV [27] and HCV [19], [22]. Here, we incorporated the inferred JM into a population genetics model, similar to the well-established Wright-Fisher model [50]. By doing so, we were able to predict the average number of generations, referred to as “escape time”, for each E2 residue to escape selective pressure (see Methods for more details). To determine escape times of these residues without the influence of the E1 protein, we utilized the E2-only model developed in our previous work [19].

Previously, the E2-only model has been shown to be capable of predicting known escape mutations from multiple E2-specific HmAbs [34], [51]–[55] (listed in Supplementary Table S2), where these mutations were shown to be associated with lower escape times compared to mutations at other residues [19] as they enable the virus to evade the associated antibody pressure. We found that this was also true for the inferred JM (*p* = 9.9 × 10^−24^; Fig. 5a), suggesting the JM to be capable of distinguishing E2 residues associated with low and high escape times.

**Fig. 5:**
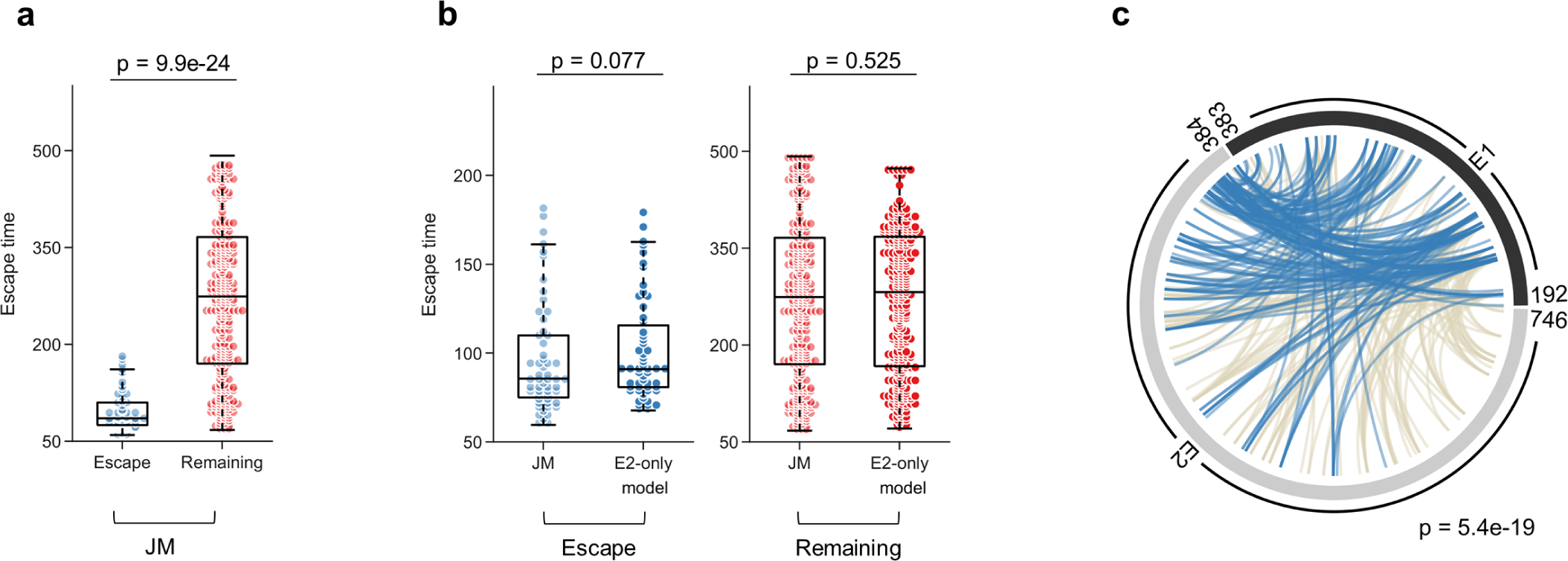
Role of E1 in facilitating viral escape from E2-specific HmAbs. (**a**) Distribution of escape times of E2 residues using the inferred JM. Residues were divided into two categories: those with known escape mutations from E2-specific HmAbs (listed in Supplementary Table S2) and the remaining E2 residues. *P*-value was calculated using the one-sided Mann-Whitney test. (**b**) Comparison of escape times of E2 residues inferred from the JM and the E2-only model for the known E2 escape mutations (left panel) and the remaining E2 residues (right panel). *P*-values were calculated using the one-sided Mann-Whitney test. (**c**) Circos plot displaying the interactions between strongly-coupled residues (Fig. 4) involving escape mutations (shown in blue) and the remaining residues (shown in beige). The reported *p*-value measures the probability of observing by a random chance at least the observed number of E2 escape mutations among strongly-coupled residues (see Methods for details).

Further comparing the escape times of E2 residues inferred from these two models, we found that the escape times of residues associated with escape mutations inferred from the JM were marginally significantly lower (*p* = 0.077; Fig. 5b, left panel) than those from the E2-only model. In contrast, there was not much difference between the escape times of the remaining E2 residues (*p* = 0.525; Fig. 5b, right panel). This suggests that the E1 protein may assist in viral escape from E2-specific antibodies. In addition, we found that the strongly-coupled inter-protein residues (Fig. 4) were statistically significantly enriched in escape mutations (*p* = 5.4 × 10^−19^; Fig. 5c). This further corroborates the potential role of E1 in mediating viral escape from neutralising antibodies.

### E. For multiple E2-specific HmAbs, E1 is predicted to provide accelerated escape dynamics

Previously, we utilized the E2-only model to assess the efficacy of each known E2-specific HmAb based on the minimum escape time predicted for its binding residues [19], [22] (see Methods for details). Our analysis above suggests that E1 may potentially assist E2 in antibody evasion, hence we further studied how this would impact the efficacy of known HmAbs predicted by the JM in comparison to the E2-only model. We first employed a binary classifier [19] to determine an optimal cut-off value (*ζ* = 96 generations) for identifying escape-resistant residues based on the JM. This binary classifier utilized known escape mutations (listed in Supplementary Table S2) as true positives and the remaining E1E2 residues as true negatives, as detailed in Methods. We subsequently evaluated each antibody by comparing the minimum escape time predicted for its binding residues with the corresponding optimal cut-off value *ζ* for each model. For this analysis, we focused on 32 HmAbs for which binding residues have been determined using global alanine scanning experiments [37], [38], [56].

Based on our previous predictions using the E2-only model, we had identified 21 E2-specific HmAbs that appear relatively easy for the virus to escape. These predictions were also consistent with the JM (Fig. 6). Among these HmAbs, studies have shown that AR1A, AR1B, AR2A, CBH-4B, CBH-4D, CBH-4G, CBH-20, CBH-21, and CBH-22 were non-neutralizing or isolate-specific [57]–[59], which further supports our predictions for both models. The remaining eight E2-specific HmAbs (212.15, 212.25, CBH-7, CBH-23, HC-1, HC33-1, HC84-20 and HCV1) were predicted to be escape-resistant by the E2-only model. However, only four (212.15 and 212.25, HC33-1, and HCV1) among these were predicted to be escape-resistant by JM (Fig. 6). The predictions of JM for these HmAbs align well with literature reports. For instance, HmAbs 212.25 and 212.15, isolated from patients who had spontaneously cleared HCV, were found to be cross-neutralizing [56]. HC33-1 and HCV1 have also been reported as potentially escape-resistant broadly neutralizing antibodies in multiple studies [37], [60]–[62]. On the other hand, of the four HmAbs (HC84-20, CBH-23, HC-1 and CBH-7) predicted to be escape-resistant by the E2-only model but not by the JM, studies have observed escape for strains isolated from patients who underwent liver transplantation for HmAbs CBH-23 and HC-1, while HmAb CBH-7 was obtained from a patient with chronic HCV infection [56], [63]. These findings suggest that E1 may play a role in facilitating HCV escape from these antibodies. Notably, the JM enabled identification of one HmAb, IGH526, that targets the E1 protein and may be escape resistant. Multiple studies have reported that IGH526 is cross-neutralizing and can target various HCV isolates from different genotypes [64], [65].

**Fig. 6:**
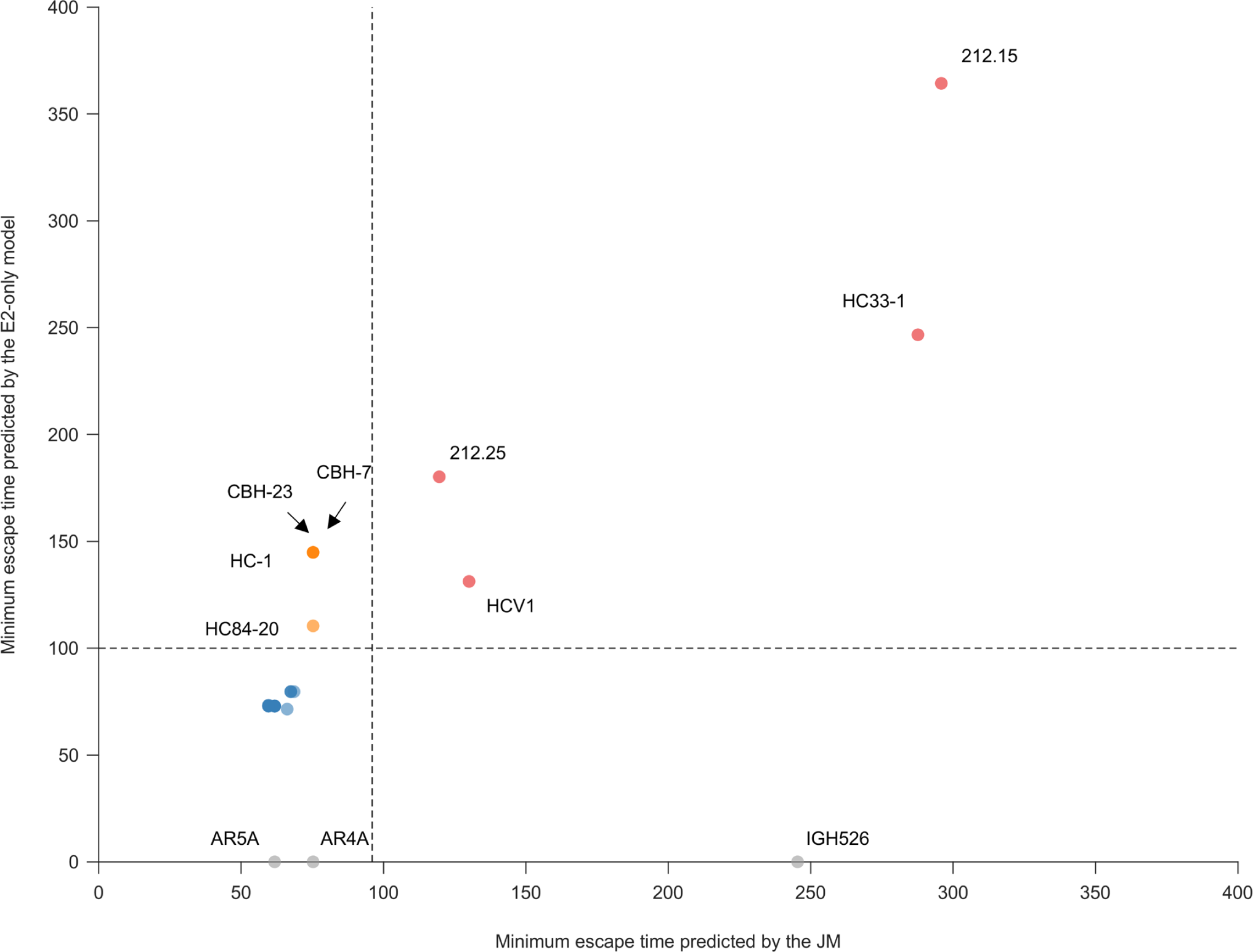
Evaluation of known HmAbs using the escape times inferred from the JM and the E2-only model. For each HmAb, escape time associated with all binding residues was predicted using both models. Each circle in the figure represents the minimum escape time associated with the binding residues of each HmAb predicted by the JM (x-axis) and the E2-only model (y-axis). Global alanine scanning mutagenesis [37], [38], [56] was used to determine the binding residues of each HmAb, where each residue of the wild-type sequence was replaced by alanine (or glycine/serine if the residue in the wild-type was alanine). We defined binding residues of each of these HmAbs as residues with relative binding (the fraction of the mutant sequence’s binding compared to the wild-type sequence) less than or equal to 20%. HmAbs predicted to be escape-resistant by both models are colored in red, the ones predicted to be escape-resistant only by the E2-only model are colored in orange, and the ones predicted to be easy to escape by both models (HmAbs 212.1.1, 212.10, A27, AR1A, AR1B, AR2A, AR3A, AR3B, AR3C, AR3D, CBH-4B, CBH-4D, CBH-4G, CBH-5, CBH-20, CBH-21, CBH-22, HC33-4, HC-11, HC84-24, HC84-26) are colored in blue. The HmAbs having binding residues in E1 (HmAbs AR4A, AR5A and IGH526) are shown in gray and plotted along the x-axis, since E2-only model could not be used to predict their escape time. The dashed line denotes the optimal cut-off value *ζ* for each model (see Methods for details).

## III. Discussion

E1 and E2 are envelope proteins of HCV that form noncovalent heterodimers. While E2 is the major target of HmAbs and a promising vaccine candidate, E1 is also important for HCV entry and assembly, and it interacts with E2. Comparing a joint model (JM) that takes into account E1E2 interactions with an independent model (IM) that does not, we have determined that these interactions are important in mediating virus infectivity and immune escape. The top E1E2 inter-protein interactions are compensatory and enriched in HVR1 and HVR2 of E2. Further using in-host evolutionary modelling, our analysis suggests that E1 may facilitate HCV in escaping E2-specific antibody responses. We have identified potentially escape-resistant HmAbs against the E1E2 complex, which could aid in the development of a robust prophylactic vaccine against HCV.

By comparing the correlation between in-vitro infectivity measurements and predictions of the JM and IM (Fig. 3), our study highlighted the importance of E1E2 inter-protein interactions in mediating viral fitness. This was further reinforced by comparing the predictions of the JM with those of a site-independent E1E2 fitness landscape model (see Methods for details), which showed that the correlation between the JM predictions and in-vitro fitness measurements was much higher than that of the site-independent model (*r* = −0.54; Supplementary Fig. S3). These findings are consistent with previous studies that have emphasized the importance of considering interactions when inferring protein fitness landscapes [19], [21], [22], [24], [25], [27], [28], [66], [67] and for identifying networks of residues that play crucial roles in protein structure and function of viruses [68]–[72].

A recent experimental study has shown that E1E2 is a fragile protein complex wherein even a single alanine mutation at 92% of positions abrogates replicative capacity of the virus [43]. Therefore, our finding that 70% of the top 300 pairs of mutations (ranked by absolute values of *J_ij_*) between E1 and E2 are compensatory suggests that these interactions may play a significant role in mediating viral fitness. To further investigate this experimentally, it would be helpful to conduct assays that quantify the change in replicative fitness by site-directed mutagenesis of the pairs of mutations identified to be associated with strong compensatory interactions (e.g., top 10) individually and simultaneously (Supplementary Data 1).

Comparing the JM and the E2-only model, we found ten residues that were predicted to be escape-resistant by the E2-only model but easy to escape according to the JM. Interestingly, four of these (residues 424, 437, 537 and 538) are known antibody binding residues, which suggests that the E1 protein may interact with these residues during antibody evasion. This motivates experimental studies for investigating the interactions between these four residues in the E2 and the E1 protein. One approach could involve longitudinal experiments [73], where the virus is allowed to infect cells in the presence of antibodies that specifically target these four residues, and changes in these residues as well as the E1 protein are monitored over time. By doing so, it could be determined if mutations arise at these residues in response to antibody pressure, and if simultaneous mutations are also observed in the E1 protein. This would provide important insights into the mechanisms by which the virus evolves to evade immune responses [74], which could ultimately inform the design of an effective vaccine against HCV.

By applying the JM and the E2-only model to evaluate the efficacy of known HmAbs, we identified 25 HmAbs with consistent predictions for both models (Fig. 6). Among these, four HmAbs were predicted to be escape-resistant, while the other 21 HmAbs were not. This motivates investigating the differences in escape dynamics [75] between these two sets of HmAbs. For instance, experimentally quantifying the average time (number of generations) it takes for the virus to escape from HmAbs 212.15, 212.25, HC33-1 or HCV1 (escape-resistant HmAbs) in comparison to HmAbs AR3A, AR3C or AR3D (non-escape-resistant HmAbs) would be a helpful follow up study.

Four HmAbs (HC-1, CBH-23, CBH-7 and HC84-20) were associated with different predictions based on the JM compared to the E2-only model (Fig. 6). We found that the different predictions for these HmAbs were due to the differences in escape times of two specific binding residues 437 and 537 by these two models, which are shared by these HmAbs. Intriguingly, these two residues are also CD81 binding residues [76]. Experiments to study the interactions between E1 and CD81 binding residues may be beneficial for discovering their potential roles in compensating viral infectivity or mediating viral entry.

## IV. Methods

### A. Inference of computational models for the E1E2 protein

To explore the role of E1E2 inter-protein interactions, we considered two types of computational models for the E1E2 protein: One taking into account the E1E2 inter-protein interactions, named the joint model (JM), and the other without the E1E2 inter-protein interactions, named the independent model (IM).

#### 1) Joint model (JM)

To infer a maximum entropy (least-biased) model for the whole E1E2 protein jointly, we downloaded 8,021 aligned E1 subtype 1a and 6,225 aligned E2 subtype 1a sequences from the HCV-GLUE database (http://hcv.glue.cvr.ac.uk) [77], [78], both with genome coverage ≥ 99%. We constructed the MSA of the whole E1E2 protein by stitching together E1 and E2 sequences based on the information in their headers, yielding 6,198 E1E2 sequences. We conducted a principal component analysis (PCA) on the pair-wise similarity matrix (6198 × 6198) of the sequences [79], where the (*i*, *j*)th entry of the similarity matrix represents the fraction of residues that are identical in sequence *i* and *j*, to remove any outlier sequences. We considered a sequence as an outlier if its corresponding value in the first PC was more than 3 scaled median absolute deviations [80] from the median of the first PC. We also excluded 264 sequences for which patients’ information was not available. After these filtering procedures, we had *M* = 5867 sequences from *W* = 871 patients. Moreover, we excluded 21 fully-conserved E1E2 residues to improve the quality of the residues. Hence, the processed MSA was composed of *M* = 5867 sequences (listed in Supplementary Data 2) and *N* = 534 residues. We constructed a least-biased maximum entropy model for the E1E2 protein that can reproduce the single and double mutant probabilities of this processed MSA (Eq. 1).

To infer parameters (**h** and **J**) of the maximum entropy model, we used the GUI realization of MPF-BML [20], an efficient inference framework introduced in [21]. This software requires an MSA as input and a vector comprising the patient weight of each sequence included in the MSA. Patient weight is computed as the inverse of the number of sequences associated with each patient. The MPF-BML parameters used for inferring the model parameters (fields **h** and couplings **J**) are: (i) *L*_1_ regularization parameters were set to 5 × 10^−4^ for both fields and couplings. (ii) *L*_2_ regularization parameters were set to 0.05 for fields, and 125 for couplings respectively and (iii) all other parameters were set to their default values. The first and second order statistics of the inferred JM matched well with those of the MSA (Fig. 1).

#### 2) Independent model (IM)

The independent model comprised of two maximum entropy models, one for the E1 protein and the other for the E2 protein. The maximum entropy models for E1 protein and E2 protein were inferred using the E1 part and the E2 part of the E1E2 processed MSA, respectively. Specifically, the MSA of both E1 and E2 consisted of *M* = 5867 sequences from *W* = 871 patients, where each sequence contains *N* = 187 residues (5 fully conserved ones were excluded) for E1 and *N* = 347 residues (16 fully conserved ones were excluded) for E2. The MSA and the patient weights were further set as the input of the MPF-BML software using the same parameters as the JM except both *L*_1_ and *L*_2_ regularization parameters were set to 50 for couplings for E1 and 15 for E2, and 5 × 10^−4^ for fields for both E1 and E2. Both the statistics of the inferred E1-only model and E2-only model lined up well with those of the respective MSAs (Supplementary Fig. S1). The final IM was a linear combination of these two models, where the energy of a full E1E2 sequence x = [x_E1_, x_E2_] is given by

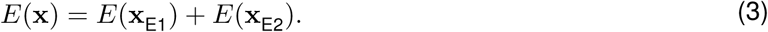

Here, *E*(x_E1_) and *E*(x_E2_) represent the energy of the E1 part x_E1_ and E2 part x_E2_ of sequence x calculated from each E1-only or E2-only model according to Eq. (1).

As we had inferred a maximum entropy E2-only model in a previous study [19], we further investigated if our previous E2-only model (inferred from 3363 sequences of E2 available at that time) was capable of capturing the statistical variations in the E2 MSA we curated in this study (5867 sequences). Our results support that this is indeed the case (Supplementary Fig. S4), suggesting that both these E2-only models are equally representative of the variations in the E2 protein sequence data. In addition, the correlation of both models with in-vitro infectivity measurements was also similar, suggesting that both E2-only models are also equally good representatives of the E2 fitness landscape (Supplementary Fig. S5).

### B. Calculation of the fraction of the correlated structure (FCS) captured by each model

FCS of the E1E2 protein complex captured by a model is given by *I*_model_*/I*. Here *I* reflects the overall strength of correlations in the protein complex [23], quantified by the difference between the site-independent model entropy (*S*_ind_) and the true entropy of the protein complex (*S*_true_). In contrast, *I*_model_ represents the strength of correlations captured by a model, calculated by the difference between the site-independent model entropy (*S*_ind_) and the inferred model entropy (*S*_model_). Below we describe how we calculated these different entropies.

*S*_ind_ was computed by considering amino acids at each E1E2 residue independently with the observed frequencies, which is given by

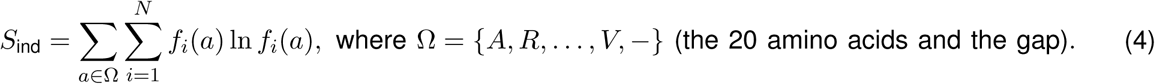

*S*_true_ was estimated using the procedure described in [23], [81] that involves incrementally sub-sampling the data and measuring its entropy. Specifically, we first randomly chose *M* sequences and calculated the the “naive estimate” of the entropy *S*_naive_(*M*) through

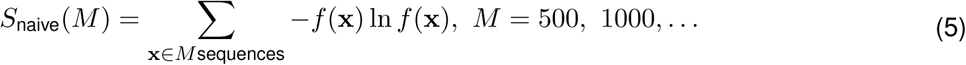

where *f* (x) is the frequency of sequence x. We repeated this procedure 100 times with different random seeds for *M* sequences (*M* = 500, 1000,...) and took the mean of *S*_naive_(*M*), denoted by ⟨*S*_naive_(*M*)⟩, over these iterations for each given *M*. As shown in [81], the naive estimate of the entropy can be well fit by

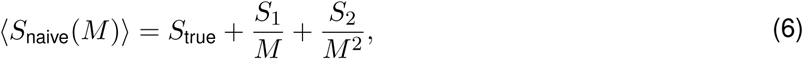

where *S*_1_ and *S*_2_ are constants that depend on the distribution of the data. They account for the bias and variance that arise due to finite sample size effects. When *M* → +∞, these correction terms vanish, and the naive estimate converges to the true entropy *S*_true_. By plotting ⟨*S*_naive_(*M*)⟩ against 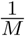, we can observe the quadratic relationship between the two variables (Supplementary Fig. S6). Extrapolating the y-intercept (when 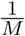 → 0) from this plot provides an estimate for *S*_true_.

We calculated *S*_model_, the entropy predicted by the inferred models, using sequence ensemble generated by a Markov Chain Monte Carlo (MCMC) procedure [23]. For the JM, the sequence ensemble comprised 99,990 full E1E2 sequences, and the model entropy was calculated as *S*_JM_ = − Σ_x_ *f* (x) ln *f* (x). For the IM, a sequence ensemble of 99,990 sequences was generated for each of the E1 and E2 protein separately using their respective individual models. The entropy for IM was calculated as *S*_IM_ = − Σ_x_*E*_1___ *f* (x_*E*_1__) ln *f* (x_*E*_1__) − Σ_x_*E*_2___ *f* (x_*E*_2__) ln *f* (x_*E*_2__), where x_*E*_1__ and x_*E*_2__ are sequences from the E1 and E2 sequence ensemble, respectively. Entropies were calculated over ten instances of MCMC runs for both JM and IM. All entropies calculated above are shown in Supplementary Fig. S7.

### C. Fitness verification

We used in-vitro experimental infectivity measurements compiled from the literature [13], [29]–[35], [37], [40]–[43] (listed in Supplementary Data 3) to investigate if our inferred models for E1E2 (JM and IM) are capable of capturing the infectivity of the virus. As experiments were conducted under different lab settings, we considered the weighted average of Spearman correlation coefficients from different experiments. This can be written as

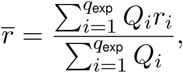

where *r_i_* is the Spearman correlation coefficient obtained from experiment *i*, and *Q_i_* is the number of measurements. *q*_exp_ is the total number of experiments.

### D. Identification of strongly-coupled residues in the E1 and E2 proteins

To identify strongly-coupled pairs of mutations (top inter-protein couplings) between the E1 and E2 proteins, we constructed “null models” to determine a threshold [67], [82]. Specifically, to maintain the observed single mutant probabilities but to break any pair-wise correlations in the E1E2 sequence data, we first constructed a “null MSA” by choosing amino acids at each residue with the observed frequencies while keeping the same number of sequences (*M* = 5867) and number of residues (*N* = 534). We then used the “null MSA” to infer a maximum entropy model, i.e, a null model. This procedure was repeated ten times, and the threshold was set as the top 0.1 percentile of the absolute mean value of *J_ij_* of these ten null models, which corresponds to roughly choosing about top 300 pairs of inter-protein couplings in the JM model. The residues that are present in these 300 inter-protein couplings are referred to as “strongly-coupled residues” throughout the manuscript.

### E. Statistical significance testing

We calculated the statistical significance of the number of strongly-coupled residues (identified by our model) in each functional regions of E1 and E2 proteins, as well as in known escape mutations (listed in Supplementary Table S2), using a *p*-value. For a given set of residues in a protein region, this *p*-value corresponds to the probability of observing at least *i* residues out of *j* strongly-coupled residues in that region, where there are *n* total residues in that protein region out of *N* total residues of a protein (187 for E1 and 347 for E2). These can be written as

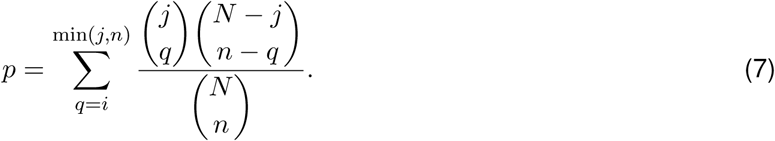

A *p*-value less than 0.05 for a protein region indicates statistically significant enrichment of residues of that region within the strongly coupled E1E2 inter-protein interactions.

### F. Visualization of interactions between strongly-coupled pairs of mutations

To visualize the interactions between strongly-coupled pairs of mutations, we utilized a Circos plot. The E1E2 residues were evenly distributed along the outer edge of the circles in Fig. 4 and Fig. 5c. The numbering of the residues was started at 192 (corresponding to the first residue of E1 according to the H77 sequence) at the 3 o’clock position and progressed in a counter-clockwise direction. Each link within the circle represents a pair of strongly-coupled mutations (ranked by the absolute values of *J_ij_* from Eq. 1).

### G. Evolutionary simulation

To quantify the average time it takes for each residue in E2 to escape with the effect of the E1 protein, we considered a viral in-host population genetics evolutionary model incorporated with the inferred JM similar to [19]. We used the “escape time” metric to represent the number of generations it takes on average for the virus with a mutation at a given residue to reach majority (frequency *>* 0.5) in a fixed-sized viral population under targeted immune pressure.

To be specific, we used a well-established Wright-Fisher model [50], where in each generation, the virus population undergoes mutation, selection, and random sampling steps. The virus population size was fixed at *M_e_* = 2000, in line with the effective HCV population size in in-host evolution [83]. For each residue *i* of the E2 protein, we formed the initial viral population with duplicates of a sequence with the consensus amino acid at residue *i*. In the mutation step, the nucleotide of each sequence was mutated randomly to another nucleotide at a fixed mutation rate *µ* = 10^−4^, consistent with the known HCV mutation rate [84], [85]. In the selection step, each sequence was selected based on its fitness predicted from the inferred JM. Specifically, we calculated the survival probability of a virus with sequence x by:

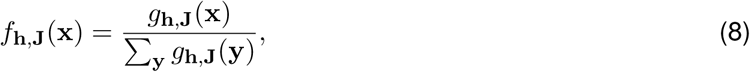

where *g***_h,J_**(x) is a function that maps the predicted energy of sequence x smoothly to a value between 0 and 1. This function is defined as:

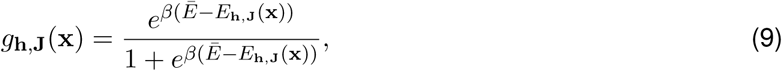

where *Ē* is the average energy of the current sequence population, while *β* ∼ 0.1 was chosen based on the slope between predicted sequence energies and in-vitro infectivity measurements [19]. To model the immune pressure at residue *i*, the fitness of all sequences having the consensus amino acid at residue *i* was decreased by a fixed value *b*, thereby providing a selective advantage to the sequences having a mutation at this residue. The value of *b* was set according to the largest value of the field parameter in the inferred landscape. Next, the subsequent generation of virus population was generated through a standard multinomial sampling process parameterized by *M_e_* and *f*_**h**,**J**_(x). This procedure was continued until the mutations at residue *i* reached a frequency *>* 0.5. The number of generations at this iteration was recorded. This process was repeated 100 times with the same initial sequence and 25 distinct initial sequences as well. The final escape time 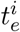 of residue *i* was the mean number of generations over all these runs of simulation.

To perform a fair comparison between the escape times predicted by the JM and those by the E2-only model in [19], we set the same simulation parameters for both models, including the fitness penalty factor *b* (10), the number of generations (500), the number of distinct sequences forming the initial population (25) for each residue, and the number of runs of simulation (100) for each distinct initial sequence. The mean escape time predicted for each residue by JM and E2-only model is provided in Supplementary Data 4.

### H. Identification of escape-resistant residues

We ran the evolutionary simulation using the JM for all E1E2 residues following the same procedure described above. We employed a binary classifier that utilized known escape mutations (listed in Supplementary Table S2) as true positives and all other residues as true negatives, which achieved an area under curve (AUC) of 0.92 (Supplementary Fig. S8a). We selected the optimal cut-off value of *ζ* ∼ 96 for determining whether a residue in the E1E2 protein is relatively escape-resistant or not based on the maximum F1 score and MCC (Supplementary Figure S8b), commonly used metrics for evaluating binary classifiers.

### I. Evaluation of the efficacy of known HmAbs

To evaluate the efficacy of known HmAbs based on the escape times obtained from the JM or the E2-only model, we adopted the following criteria. We compared the minimum escape time 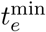 predicted for a HmAb’s binding residues [37], [38], [56] with the cut-off value (*ζ*) for each model. If 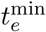 of a HmAb was greater than *ζ* for a model, that HmAb was characterized as relatively escape-resistant by that model, and vice versa.

### J. Site-independent model

In order to compare JM with a model that ignores all interactions between residues, we defined a site-independent E1E2 fitness landscape model that is characterized solely by the “fields” h as follows

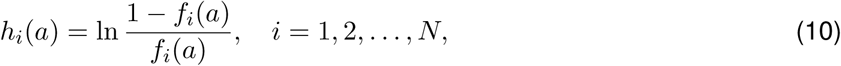

where *f_i_*(*a*) is the frequency of observing amino acid *a* at residue *i*.

## Data and code availability

- All data used in this work is publicly available. Top 300 pairs of inter-protein couplings obtained from the JM are listed in Supplementary Data 1. Accession numbers of E1E2 sequences used for inferring JM and IM are listed in Supplementary Data 2. The E1E2 infectivity measurements, used for correlating with predictions obtained from the inferred JM and IM, are included in Supplementary Data 3. The mean escape time predicted for each residue by JM and E2-only model is provided in Supplementary Data 4.
- The GUI-based software implementation of the MPF-BML method [21], used for inferring the fitness landscape model, is available at https://github.com/ahmedaq/MPF-BML-GUI [20]. Data and scripts for reproducing the results of this manuscript are available at https://github.com/hangzhangust/HCVE1E2.
- Any additional information related to the data reported in this paper is available from the lead contact upon request.

## Acknowledgements

H.Z. and A.A.Q. were supported by the Hong Kong Research Grants Council (grant number 16204519). A.A.Q. and M.R.M. were supported by the Australian Research Council through Discovery Project (DP 230102850). A.A.Q., R.A.B., and M.R.M. were supported by Australia’s National Health and Medical Research Council (NHMRC) through IDEAS Project (2020192). M.R.M. is the recipient of an Australian Research Council (ARC) Future Fellowship (project number FT200100928). R.A.B. is a fellow funded by NHMRC.

## Supplementary Information

### HCV E1 influences the fitness landscape of E2 and may enhance escape from E2-specific antibodies

#### Supplementary Figures

**Fig. S1:**
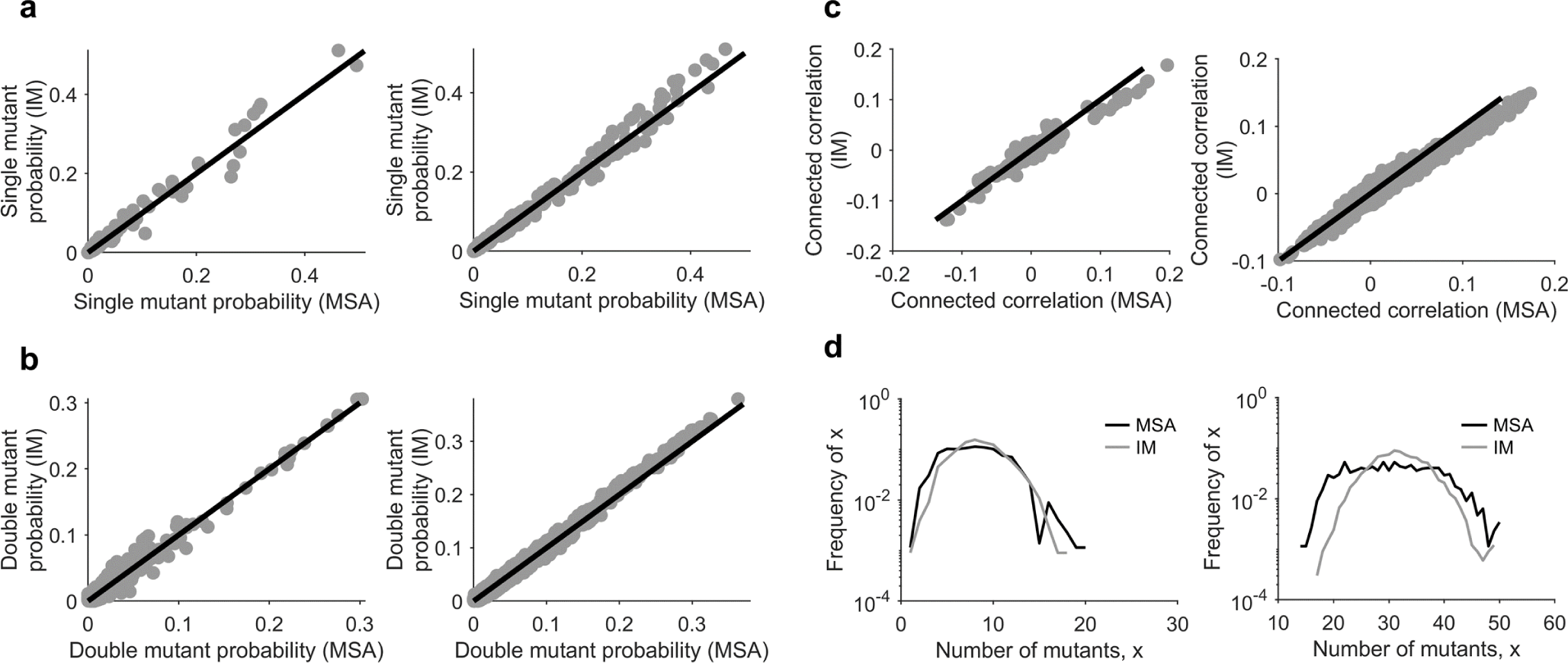
Statistical validation of the inferred IM for the E1E2 protein. Comparison of the (**a**) single mutant probabilities, (**b**) double mutant probabilities, (**c**) connected correlations, and (**d**) distribution of the number of mutants per sequence obtained from the MSA (E1 in left panels and E2 in right panels of each subfigure) and those predicted by the inferred IM. Samples were generated from the inferred model using the MCMC method [1].

**Fig. S2:**
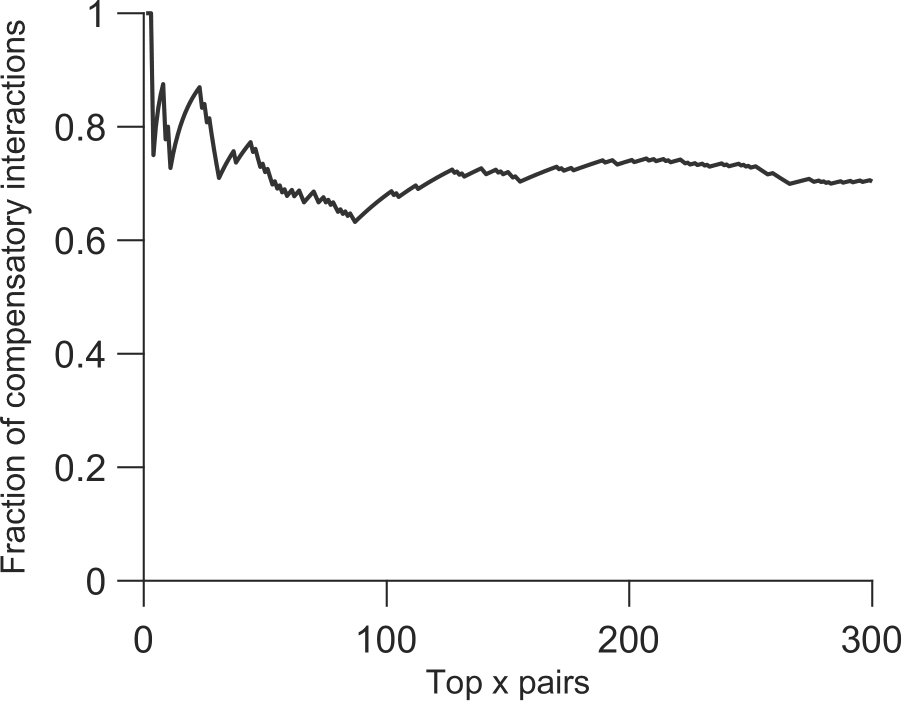
Robustness of the fraction of compensatory E1E2 inter-protein interactions (Fig. 4) to the number of top inter-protein couplings selected. Top pairs of E1E2 inter-protein couplings were ranked by the absolute values of *J_ij_* from Eq. 1. Fraction of compensatory interactions were calculated by the number of negative values of *J_ij_* divided by top x pairs of inter-protein couplings considered.

**Fig. S3:**
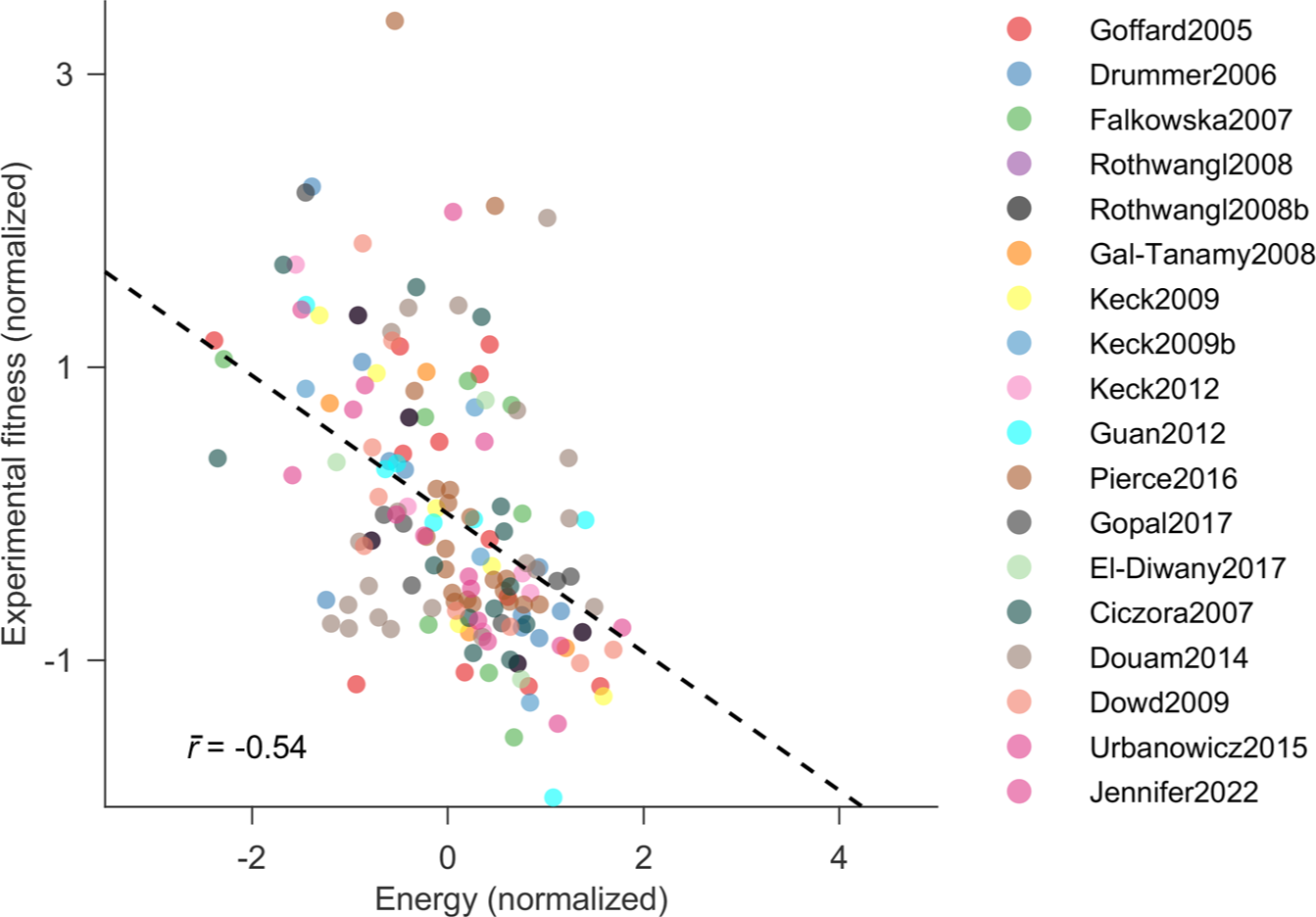
Correlation between infectivity measurements and predictions obtained from a site-independent model. The site-independent model was inferred using only single mutant probabilities (see Methods for details). Legend shows the references from which fitness measurements were compiled [2]– [17].

**Fig. S4:**
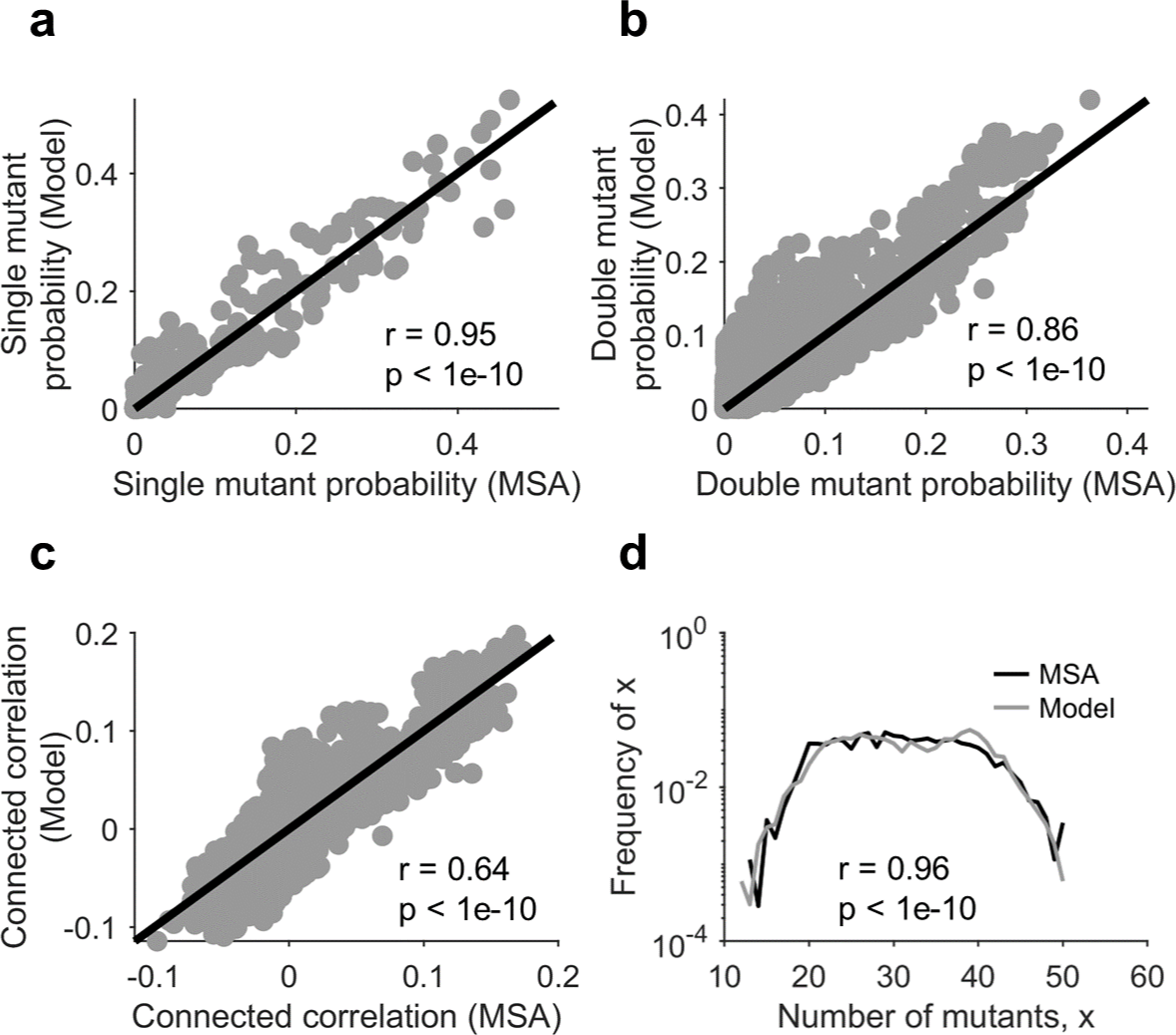
E2-only model inferred in our previous study [18] reproduces statistics of the MSA based on the latest E2 sequence data. The model in [18] was inferred using HCV E2 sequence data up until September 2017. Comparison of the (**a**) single mutant probabilities, (**b**) double mutant probabilities, (**c**) connected correlations, and (**d**) distribution of the number of mutants per sequence obtained from the MSA and those predicted by the previous E2-only model. Samples were generated from the previous E2-only model using the MCMC method [1]. Pearson correlation coefficient and the associated *p*-value is shown for each subfigure.

**Fig. S5:**
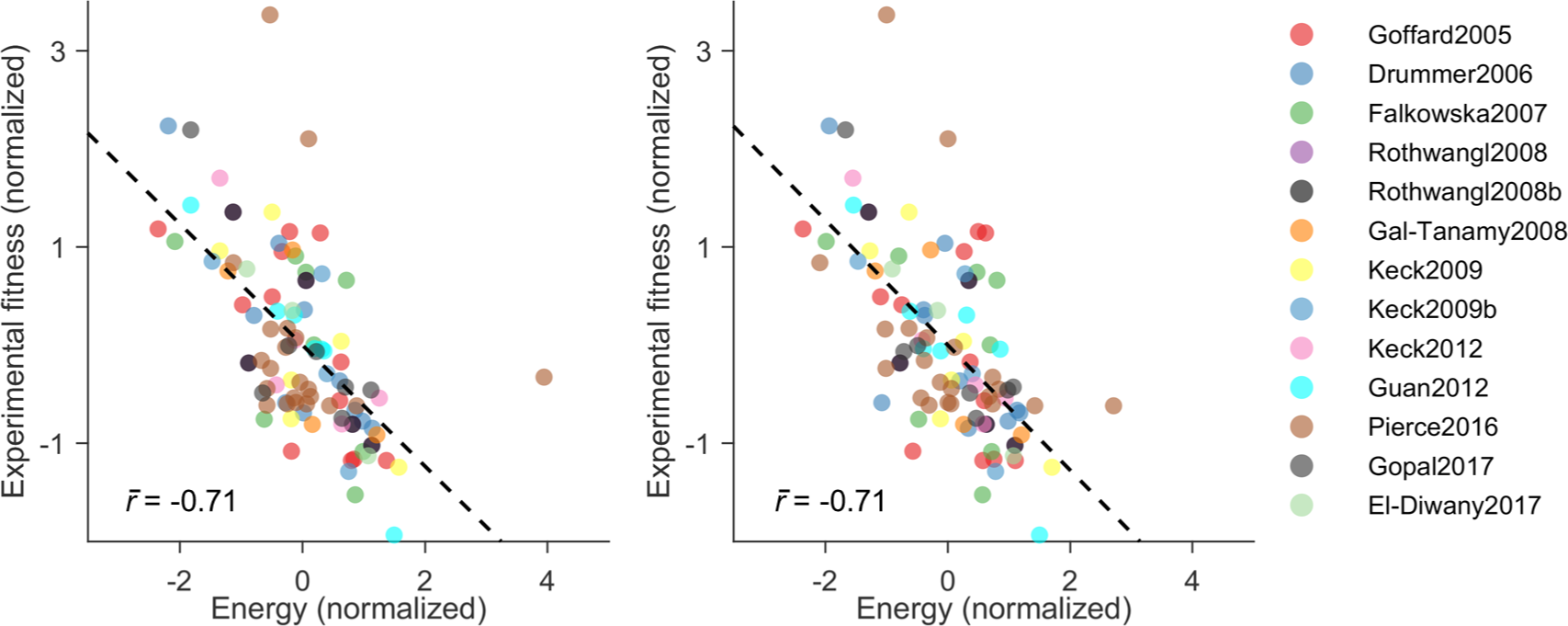
Comparison of the fitness prediction of the E2-only model inferred in this work (left panel) and in our previous study [18] (right panel). The model in [18] was inferred using HCV E2 sequence data up until September 2017. Normalized energies computed from both models correlate strongly with E2-only experimental fitness measurements. Legend shows the references from which E2-only fitness measurements were compiled [2]–[12].

**Fig. S6:**
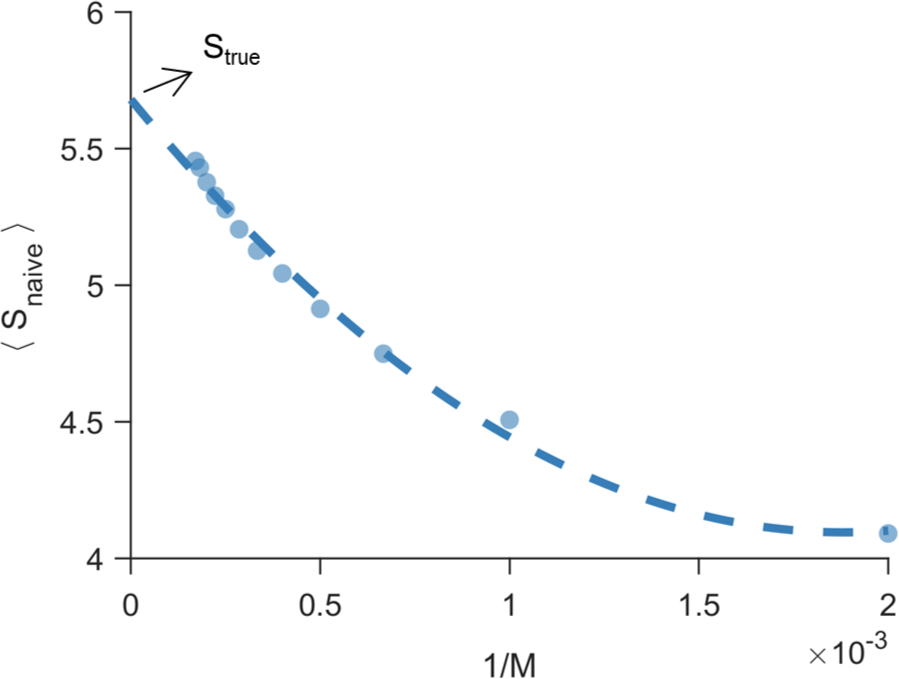
Estimate of the true entropy. ⟨*S*_naive_(*M*)⟩ vs. 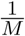 (shown as circles) can be well fit by 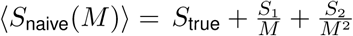 (shown as dashed line), where *S*_1_ and *S*_2_ are constants [19], [20]. The y-intercept of the fit provides an estimate for the true entropy, *S*_true_ (see Methods for details).

**Fig. S7:**
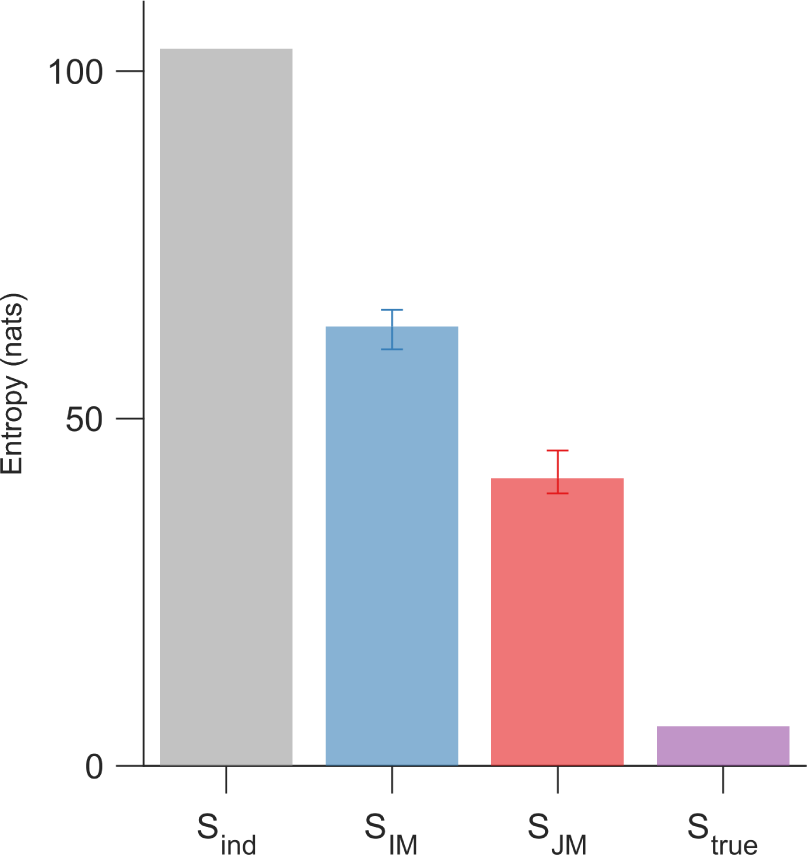
Entropy calculated from different models for the E1E2 protein. *S*_ind_ denotes the entropy of a site-independent model of E1E2 protein, where we considered choosing amino acids at each residue of the E1E2 protein independently with the observed frequencies. *S*_IM_ and *S*_JM_ are entropies calculated from the inferred IM and the JM (calculated over 10 instances of MCMC runs), and *S*_true_ is the estimated true entropy (see Methods for details).

**Fig. S8:**
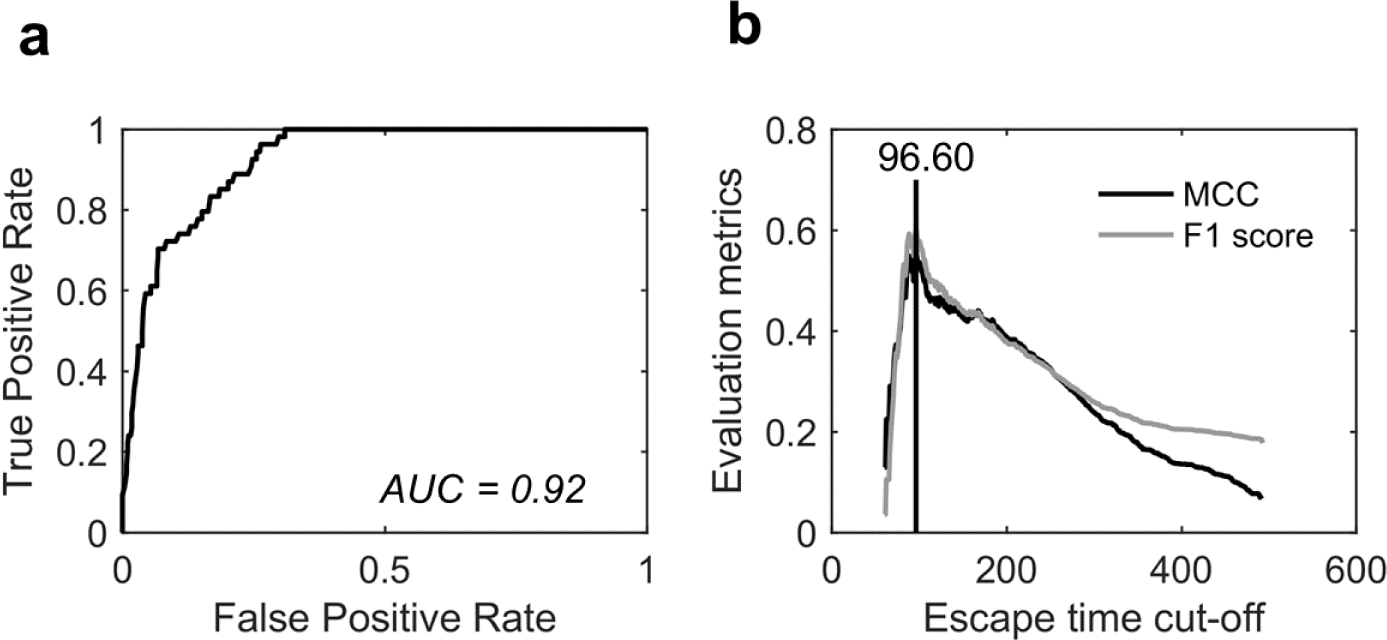
Binary classifier designed to determine the optimal cut-off for escape time based on experimentally or clinically identified escape mutations. The classifier used a Receiver Operating Characteristic (ROC) curve to identify known E2 escape mutations (listed in Supplementary Table S2) using the escape time metric. The optimal cut-off value was determined by maximizing the F1 score and the Matthews Correlation Coefficient (MCC). In this classification, E2 residues with known escape mutations were considered as true positives, while all remaining residues were considered true negatives.

#### Supplementary Tables

**TABLE S1:** List of functional regions of E1 and E2 proteins along with the statistical significance of enrichment of strongly-coupled residues in each region.

| **Regions of E1** | ***p*-value** | **Regions of E2** | ***p*-value** |
| --- | --- | --- | --- |
| Putative fusion peptides of E1<br>(Fusion E1, residue 265-296) | 0.7736 | Hypervariable region 1<br>(HVR1, residue 384-410) | 7.5e-8 |
| Stem region of E1<br>(Stem E1, residue 309-349) | 0.8251 | Hypervariable region 2<br>(HVR2, residue 460-482) | 3.0e-4 |
| Transmembrane domain of E1<br>(TMD E1, residue 350-381) | 0.5719 | Intergenotypic variable region<br>(igVR, residue 570-580) | 0.5848 |
| N-terminal domain of E1<br>(NTD E1, residue 192-239) | 0.7233 | Stem region of E2<br>(Stem E2, residue 662-717) | 0.9991 |
| Other regions of E1<br>(Other E1) | 0.0776 | Transmembrane domain of E2<br>(TMD E2, residue 718-742) | 0.9961 |
|  |  | Other regions of E2<br>(Other E2) | 0.9858 |

**TABLE S2:** List of known escape mutations from E2-specific HmAbs.

| **Escape residues** | **HmAbs** | **Reference** |
| --- | --- | --- |
| 408 | HC33-4 | [21] |
| 384, 386, 388, 390, 391, 393, 394,<br>395, 396, 397, 398, 399, 400, 401,<br>402, 403, 404, 405, 407, 410 | HmAbs targeting HVR1 | [22] |
| 431 | CBH-2 | [23] |
| 431, 435, 444, 446, 466, 482, 501,<br>528, 531, 538, 580, 610, 636, 713 | CBH-8C, CBH-2, CBH-5, HC-2,<br>HC-11 | [7] |
| 391, 394, 401, 415, 417, 434, 444,<br>608 | HCV1 | [24] |
| 416, 422, 424, 431, 433, 438, 442,<br>446, 453, 456, 461, 475, 482, 520,<br>524, 531, 533, 557, 558, 560 | CBH-2, CBH-5, HC84-22, HC84-26,<br>AR3A, AR3B, AR3C, AR3D | [25] |
| 431, 438, 442 | AR3A | [26] |

